# Flower bud cooling protects pollen development and improves fertility during heatwaves

**DOI:** 10.1101/2024.08.18.608441

**Authors:** Martijn J. Jansen, Stuart Y. Jansma, Klaske M. Kuipers, Wim H. Vriezen, Frank F. Millenaar, Teresa Montoro, Carolien G.F. de Kovel, Fred A. van Eeuwijk, Eric J.W. Visser, Ivo Rieu

## Abstract

Early pollen development is a bottleneck for plant fertility in heatwave conditions, thus affecting yield stability. Mechanisms that protect this process and explain variation in tolerance level between genotypes are poorly understood. Here we show that sepal transpiration in young, still closed, flower buds reduces the impact of heat on developing tomato pollen and that this mechanism is enhanced by the major tomato pollen thermotolerance QTL, qPV11. By direct measurement of the flower bud core temperature and transpiration we show this process, which we term ‘flower bud cooling’, depends on heat-induced opening of sepal stomata and that the transpiration enhancing effect of qPV11 requires functional stomatal regulation and is specific to the sepals. Large-scale evaluation of populations in both a production field and greenhouse showed that qPV11 improves pollen viability and fruit set in heatwave-affected complex cultivation environments. These findings highlight enhanced flower bud cooling as a naturally evolved protection mechanism against heatwaves and qPV11 as genetic component in the differential regulation of transpiration between reproductive and vegetative tissues and candidate variant for the breeding of climate-resilient tomato cultivars.

---

The intensity, frequency, and duration of heatwaves continues to increase since the mid-20th century due to ongoing climate change, leading to an expected quadrupling of the areas impacted by such heat extremes by 2040 compared to the first decade of this century^1,2^. Reproductive development is the most heat sensitive process in the plant life cycle, resulting in a strong negative effect of heatwaves on reproductive crop yields and food production security and environmental footprint^3,4^.

Male reproductive development is particularly sensitive to long-term, mildly elevated temperatures that characterise a heatwave and pollen death is a major contributor to heatwave-induced loss of seed and fruit set^5–7^. This type of temperature profile is especially harmful during early pollen development with a time-window of sensitivity around microsporogenesis^8,9^. Previous studies have shown that the impact of heatwaves on pollen development is independent of its effect on the rest of the plant, suggesting that the temperature of the flower bud itself is crucial for pollen viability^9–15^.

The temperature of flower buds can deviate significantly from the ambient air temperature. Sub-ambient temperatures have been observed under mild heat in flowers or flower buds of rice^16^, wheat^10^, soy^17^, groundnut^18^, pea^19^ and several non-crop species^20,21^. However, the impact on reproductive output and the underlying mechanisms remain largely unexplored. Recent research suggests that the lower temperature of mature common bean flowers results from transpirational cooling across the entire canopy^22^. In contrast, in cleistogamous soybean, reduced internal temperature of mature flowers was achieved by local transpiration from the sepals^17^. Similar reduced flower bud temperatures and the presence of sepal transpiration were found under extreme heat in the later stages of tomato flower bud development^23^. As microsporogenesis is the reproductive process that is most sensitive to heatwave-like temperature profiles, it is a compelling question whether active cooling occurs in young flower buds, too, and provides protection to pollen development.

Tomato (*Solanum lycopersicum*) is one of the major vegetable crops globally, and its production is strongly affected by climate change^24^. Exposure to a heatwave during reproductive development can significantly reduce pollen numbers and viability, leading to a decrease in fruit set^5–7^. During early flower bud development, the reproductive organs are closely encapsulated by sepals containing stomata, which provides the potential for transpiration-mediated flower bud cooling^9,25,26^.

This study demonstrates the existence of flower bud cooling during early microspore development in tomato, its dependency on sepal transpiration and stomatal regulation, and its role in pollen thermotolerance. Additionally, we show that the pollen thermotolerance QTL qPV11 enhances transpirational flower bud cooling and thereby improves reproductive thermotolerance both under greenhouse and open field conditions.

## Main

### Under high air temperatures, tomato flower buds maintain a sub-ambient temperature

To assess the ability of tomato plants to cool their flower buds during microsporogenesis, we simultaneously measured the temperature of the core of microspore-stage buds and the air (Extended data Fig. 1). During the initial heating phase of a heatwave-like event, flower bud temperature closely followed the increasing air temperature, including the climate cabinet induced oscillations, indicating a high thermal conductivity of the plant tissue (Fig. 1a). However, after approximately 30 minutes, the flower bud temperature began to decrease even as the air temperature continued to rise to ∼34 °C.

**Figure 1:**
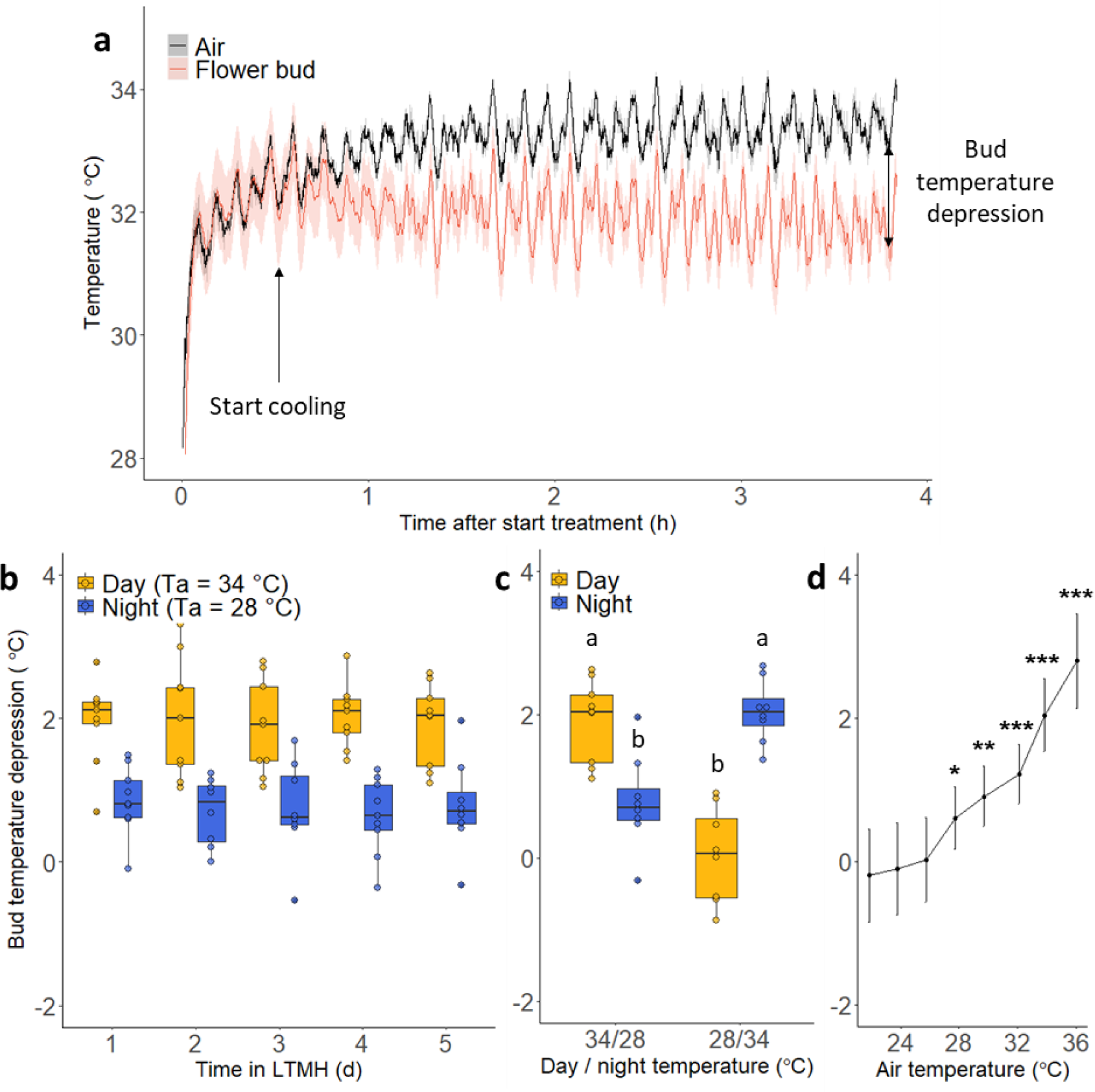
Flower buds are cooled under high temperatures. (a) Tomato flower bud temperature (red) and air temperature (black) during the initial phase of a heatwave-like temperature profile. Indicated are mean temperatures and 95% confidence intervals, n = 2 air sensors, n = 9 plants. Arrows indicate the moment the bud temperature starts to deviate from the air temperature and the stabilized bud temperature depression (BTD). (b) Over 5 days of LTMH, the bud temperature depression is constant during both hot days (34 °C) and milder night (28 °C). Indicated are median and interquartile ranges of mean BTD during day (9:00 - 18:00) and night (21:00-07:00), n = 8-9 plants. (c) Bud temperature depression during a hot day and mild night (34/28 °C) and hot night and mild day (28/34 °C) regime. Indicated are median and interquartile ranges of mean BTD, n = 8-9 plants. Letters indicate groups with significantly different mean BTD (two-way ANOVA, Tukey’s post-hoc test, P<0.05). (d) The level of bud temperature depression is dependent on the air temperature with stronger cooling occurring at higher air temperatures. Indicated are mean BTD and standard deviations, n = 8 plants. Deviation of BTDs from 0 was determined by one-tailed one-sample t-test with Benjamini and Hochberg correction (*P = 0.005, **P = 0.0005, ***P<0.0001).

This difference between flower bud and air temperature, referred to as the flower bud temperature depression (BTD), increased over the next 30 minutes and then stabilized at 2 °C (Extended data Fig. 2). The average temperature depression remained consistent during recurrent warm days and was lower during the cooler nights (Fig. 1b). Interestingly, flower bud cooling occurred independently of light and circadian rhythm, as evidenced by similar levels of cooling at high temperature, when day and night temperatures were interchanged (Fig. 1c). To determine whether the extent of flower bud temperature depression depends on the ambient air temperature, we measured the flower bud temperature at air temperatures from 22 to 36 °C, maintaining a constant air vapor pressure deficit (VPD) and radiation. At temperatures below 26 °C the flower bud and air temperatures were similar, whereas at temperatures above 26 °C, flower buds were cooler compared to the environment, with the bud temperature depression increasing with higher air temperature (Fig. 1d). The increased cooling, however, was not sufficient to fully offset the rising air temperatures, which is consistent with the increasing damage to pollen development found at increasing air temperatures. Interestingly, the changing point at which cooling was induced is similar to that in the pea apical bud^19^ (26 °C), but substantially lower than in rice spikelets (around 30 °C)^16^, groundnut flower buds (between 28 and 34 °C)^18^ and leaves of many species (30 °C)^27^. These differences in changing point are consistent with the (agro)ecological temperature thresholds of these crops^28,29^. Given that even minor differences in temperature during the sensitive phase strongly impact pollen viability and fruit set under LTMH conditions^3,6^ and that LTMH acts locally at the level of the flower bud^9^, flower bud cooling evolved likely to mitigate the impact of heat on reproduction.

**Figure 2:**
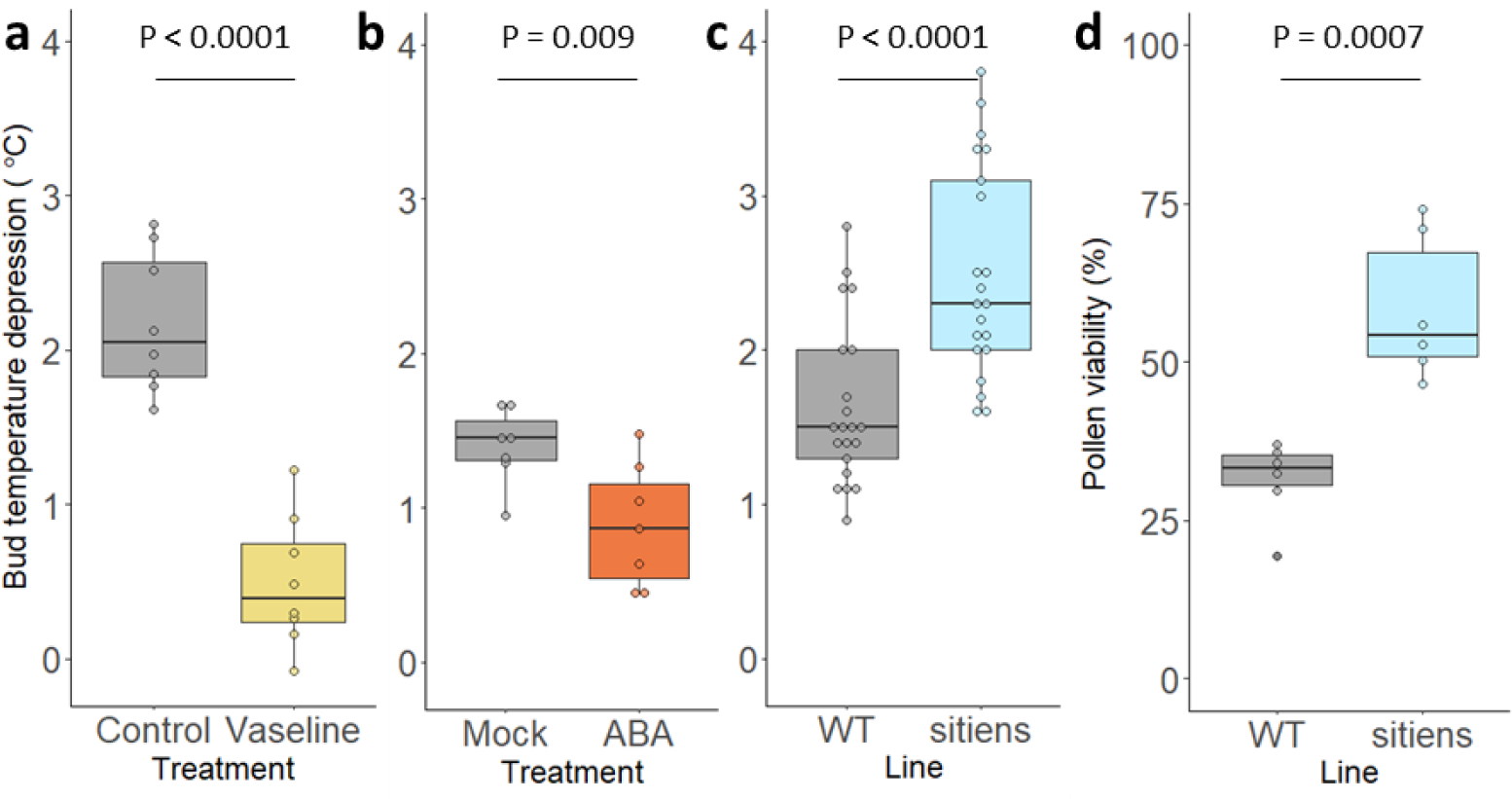
Manipulation of the gas exchange capacity of flower buds impacts the bud cooling capability under LTMH and pollen heat tolerance. (a,b) Bud temperature depression under LTMH of flower buds with reduced local gas exchange due to application of Vaseline on the flower buds (a) or treatment with ABA (b). (c,d) Bud temperature depression and pollen viability under LTMH of the ABA-deficient mutant sitiens with constitutively open stomata. Indicated are median and interquartile ranges of plant mean BTD and pollen viability on the 5^th^ day and in the 3^rd^ week of LTMH, respectively (a, n = 8 plants, b, n = 7 plants, c: n = 21 flower buds, d: n = 6 plants). Significant differences were determined by respectively a two-sided paired samples t-test, two-way ANOVA including block effects, Mann-Whitney test and two-sided t-test.

### Flower bud cooling is due to active regulation of sepal stomata

The temperature of plant tissue is determined by a balance between net incoming radiation, conduction, convection and evapotranspiration^30^. At high ambient temperatures, plants rely especially on evaporative cooling to prevent heat damage to leaves^31^. The application of the energy balance model for plant tissue thermoregulation showed that the observed flower bud temperatures at air temperatures above 26 °C can only be achieved if the vapor resistance of the sepal surface is reduced through stomatal opening (Extended data Fig. 3). Indeed, we found that reducing stomatal conductance, either by applying wax to the sepals covering the bud or through treatment with the plant hormone abscisic acid (ABA), significantly impaired flower bud cooling (Fig. 2a,b). This indicates that vapor exchange and open stomata are required for flower bud cooling in tomato, similar to mature flower cooling in cleistogamous soy^17^. This is further supported by the observed delay and subsequent concave response curve of the bud temperature depression during the onset of LTMH (Fig. 1a), which closely mirrors the described increase in stomatal aperture and vapor exchange under high light conditions^32,33^. In contrast, transpiration responds quickly to changing VPD if stomatal aperture is constant^31^.

**Figure 3:**
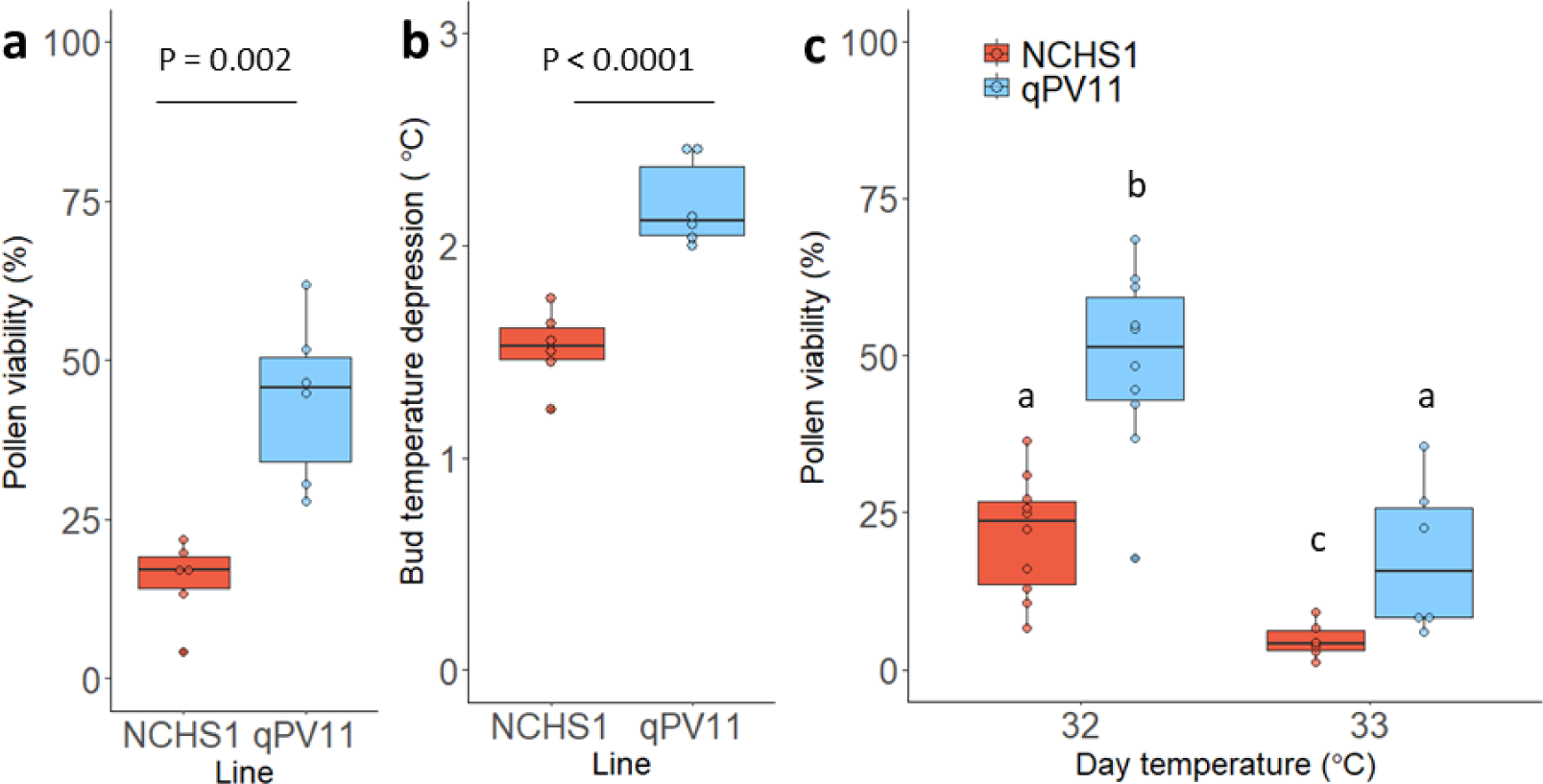
The pollen thermotolerance QTL qPV11 increased pollen viabilities under LTMH by enhanced flower bud cooling. (a,b) The effect of qPV11 on the pollen viability after LTMH (a) and bud temperature depression during LTMH (b). Indicated are median and interquartile ranges of mean pollen viability and bud temperature depression of a NIL containing qPV11 and its respective background NCHS-1 (n = 6 plants). Significant differences were determined by two-way ANOVA including block effects. (c) The effect of an increase in temperature on pollen viability in the NIL with qPV11. The effect of a difference in flower bud temperature was mimicked by growing NCHS-1 and the qPV11 NIL under standard LTMH conditions, with a day temperature of 32 °C, and under LTMH with a day temperature of 1 °C higher. Indicated are median and interquartile ranges of mean pollen viability under LTMH with a day temperature of 32 °C (n = 10 plants) and 33 °C (n = 6 plants). Letters indicate groups with significantly different mean pollen viabilities (two-way ANOVA, Tukey’s post-hoc test. P<0.01).

Under LTMH conditions, the ABA-deficient tomato mutant *sitiens*, which has constitutively open stomata^34^, exhibited a significantly higher degree of bud temperature depression than the wild type (Fig. 2c), implying that a further increase in cooling is possible. Notably, we found that *sitiens* produced more viable pollen when grown under LTMH conditions (Fig. 2d), thus suggesting enhancement of transpiration and flower bud cooling enhances male reproductive heat tolerance.

### The pollen thermotolerance QTL qPV11 acts by enhancing flower bud cooling

Extensive screening of commercial cultivars and wild relatives of crops, has led to the identification of natural variation in plants’ abilities to withstand high temperatures^35,36^. This natural variation has been utilized to identify several quantitative trait loci (QTL) associated with reproductive heat tolerance, including qPV11, a major QTL in tomato for pollen viability under LTMH ^37^. This QTL was introduced into a distinct tomato line, NCHS-1, to create a near isogenic line (NIL) that confirmed the impact of qPV11 on pollen thermotolerance (Fig 3a). Notably, at an ambient temperature of 32 °C, the presence of qPV11 resulted in an additional decrease in flower bud temperature of about 0.7 °C (Fig 3b). At an ambient temperature of 33 °C, plants with qPV11 showed a similar degree of pollen damage to plants without qPV11 at 32°C (Fig 3c). Therefore, qPV11 increases the pollen thermotolerance of tomato plants by improving the cooling capacity of young flower buds. This is supported by the fact that the effect of qPV11 on bud temperature depression and pollen thermotolerance was very similar to that of the *sitiens* mutation (Fig 2c,d).

### qPV11 improves flower bud cooling by affecting sepal stomatal regulation

The cooling of flower buds depends on transpiration. To measure the effect of qPV11, we compared the transpiration rate of NCHS-1 flower buds with those of the NIL at 32 °C. The result shows that qPV11 increased the transpiration rate of flower buds, both per sepal surface area and per total bud weight (Fig. 4a,b). To test whether the positive effect of qPV11 on transpiration was due to increased stomatal opening, we tested the impact of qPV11 on transpiration rate, while inhibiting normal stomatal regulation. Both, constitutive opening of sepal stomata, through application of fusicoccin, and constitutive closure of sepal stomata, through application of ABA, eliminated the positive effect of qPV11 on transpiration rate (Fig. 4b). This suggests that qPV11 acts through regulation of stomatal responses. Interestingly, qPV11 appears specific to flowers as we did not measure any impact of the QTL on transpiration rate or temperature depression of the leaves (Fig. 4c, Extended data Fig. 4).

**Figure 4:**
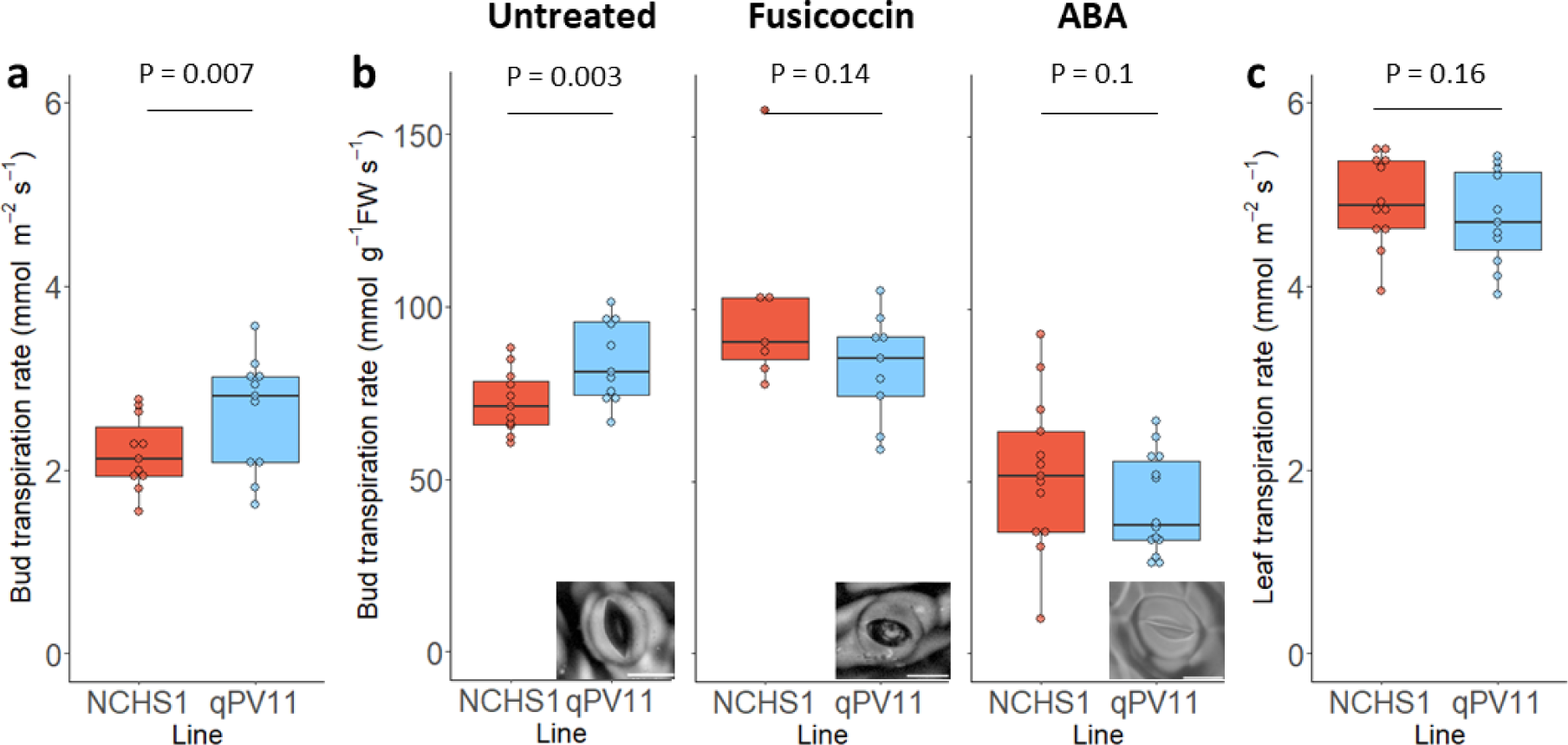
The effect of qPV11 on flower bud transpiration under LTMH depends on sepal stomatal regulation. The effect of introgression of qPV11 on bud and leaf transpiration rate per sepal surface area (a,c) and on bud transpiration rate per gram flower bud with disrupted stomatal responses (b) on the 5^th^ day of LTMH. Stomatal responses were disrupted by treatment with fusicoccin or ABA leading to respectively fully open or closed stomata (representative sepal stomatal prints from plants with qPV11 included in bottom right, Scale bars: 20 µm). Indicated are median and interquartile ranges of mean bud transpiration rate per sepal surface area (a) and bud transpiration rate per gram fresh bud weight (b) of a NIL containing qPV11 and its respective background NCHS-1. n = 11, n = 11, n = 7-9, n = 13-14 and n = 11 plants for each respective measurement and treatment Significant differences were determined by two-way ANOVA including block effects.

### qPV11 improves thermotolerance in actual crop production environments

The applicability of traits and QTLs for crop climate adaptation depends on their genetic and environmental stability. To test this, we created a processing tomato RIL population from four different tomato lines, including Nagcarlang, which contained qPV11 and cultivated the F6 population in a commercial production field in Extremadura, Spain. An analysis of pollen viability and fruit set after a natural heatwave revealed that the presence of qPV11 resulted in increased pollen thermotolerance and end-of-season fruit set (Fig. 5a,b). Additionally, a fresh market tomato RIL population was created from four tomato lines, including Nagcarlang, and grown in a commercial production greenhouse in Almeria, Spain. Phenotyping of the F4 population after the occurrence of a strong heatwave confirmed the positive effect of qPV11 on pollen viability and fruit set under more severe heat (Fig. 5c,d). These findings show that enhanced flower bud cooling by qPV11 improves reproductive heat tolerance even in complex production environments.

**Figure 5:**
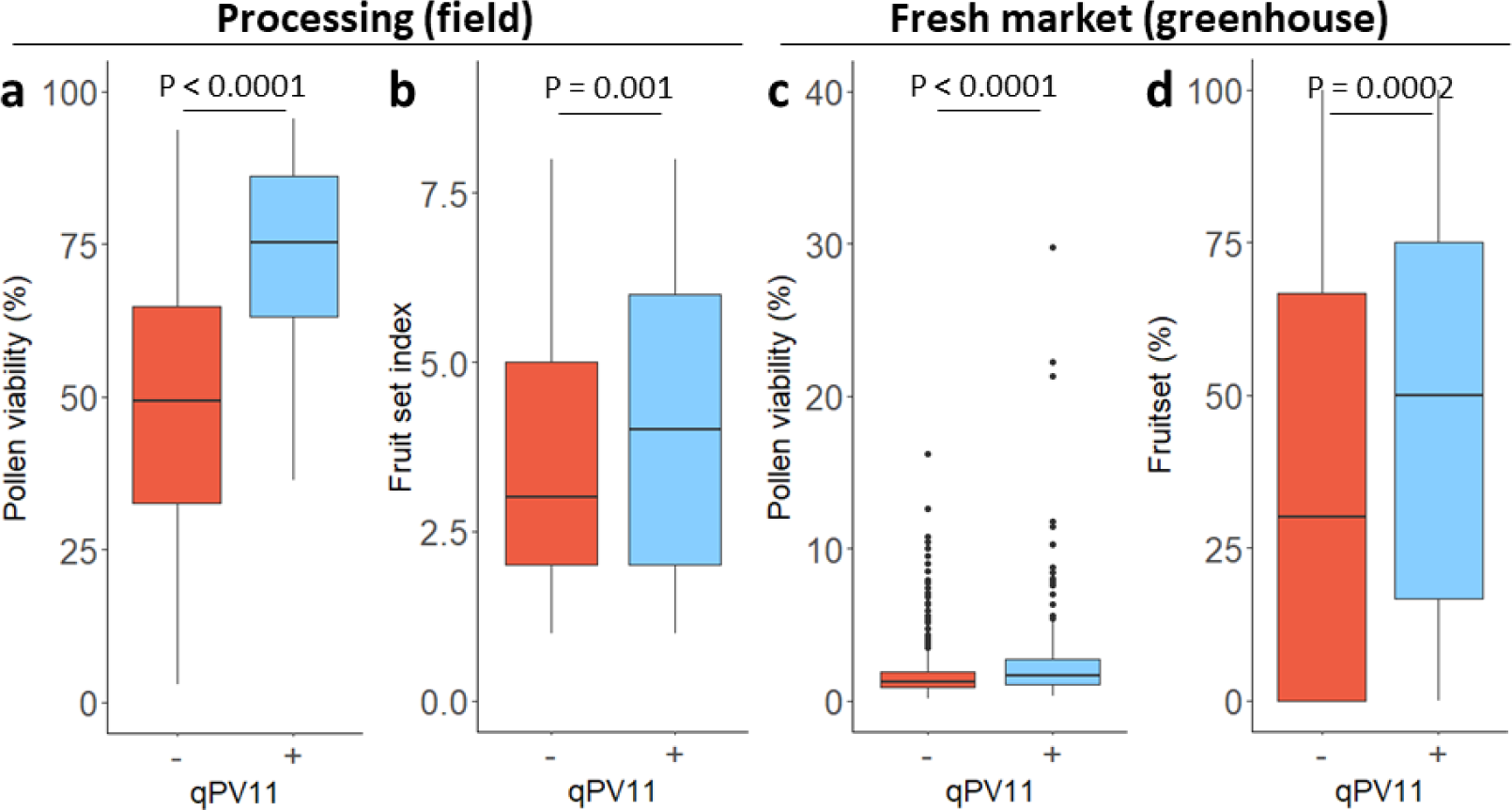
The effect of qPV11 on pollen viability and fruit set in the field and greenhouse. The effect of qPV11 on pollen viability of flowers of which microsporogenesis occurred during a heatwave (a) and end of season fruit set (b) of determinate processing-type RILs grown in a production field during a hot season (n = 109 qPV11+ and 539 qPV11-plants). The effect of qPV11 on pollen viability (c, n = 201 qPV11+ and 515 qPV11-plants) and fruit set (d, n = 171 qPV11+ and 438 qPV11-plants) of flowers of which microsporogenesis occurred during an extreme heatwave in an indeterminate, fresh-market-type RIL population grown in a production greenhouse. Plants are separated on the homozygous presence of the Nagcarlang allele (blue) or contrasting allele (red) for qPV11. Indicated are median and interquartile ranges of mean pollen viability, fruit set breeding score and mean seeded fruit set per flower. Significant differences in pollen viability were determined by two-tailed Welch t-test and in fruit set by Mann-Whitney test.

## Discussion

Climate change is causing more frequent and intense heatwaves, which in turn lead to increased occurrence of reproductive heat stress and crop yield losses worldwide. Plant microsporogenesis is a particularly heat-sensitive process^9^. We showed that at high temperature, young tomato flower buds are cooled by means of sepal transpiration and that natural variation in pollen thermotolerance encoded by qPV11 is due to further enhancement of sepal transpiration, through stomatal regulation. This trait improves yield stability of tomato crop exposed to a heatwave in actual production environments.

The enclosure of young flower buds in stomata-bearing sepals is widespread in the plant kingdom^38^, as is heat-induced stomatal opening^39–42^. Therefore, it is likely that flower bud cooling also exists in other species besides tomato, offering new strategies for improving reproductive heat tolerance in horti- and agricultural crops. It remains to be tested whether enhanced flower bud cooling can be combined with cellular heat tolerance traits, e.g. based on manipulation of pollen flavonol levels^43^, to further raise the heat tolerance level, and whether reduction of the flower bud temperature also mitigates the impact of heat on female reproductive development^44^.

Interestingly, qPV11 affects transpiration of the sepals and not the leaves. This genetic uncoupling is analogous to the uncoupling of stomatal behaviour of sepals and leaves seen when heat exposure is combined with drought^17,23^. The ability to improve cooling of reproductive tissues without increasing leaf transpiration allows plants to protect sensitive reproductive processes while having a limited impact on overall water use efficiency.

Heat-induced stomatal opening occurs guard-cell autonomous, independent of CO_2_ assimilation status and light condition^45,46^, consistent with our findings of bud temperature depression during hot nights. At least three signalling pathways have been implicated in heat-induced stomatal opening: a phototropin-dependent pathway^46^, a ROS-dependent pathway requiring RBOHD activity^47^ and a pathway leading to reduced ABA sensitivity^45^. Under a combination of heat and drought differential stomatal behaviour has been attributed to a flower specific increase in the ABA metabolism^17,23^. Identification of the causal genetic variation in qPV11 will shed light on the mechanism behind organ specific heat-induced stomatal regulation and suggest additional targets and approaches to enhance reproductive heat tolerance and improve climate resilience and sustainability of crops.

## Extended data

**Extended data figure 1:**
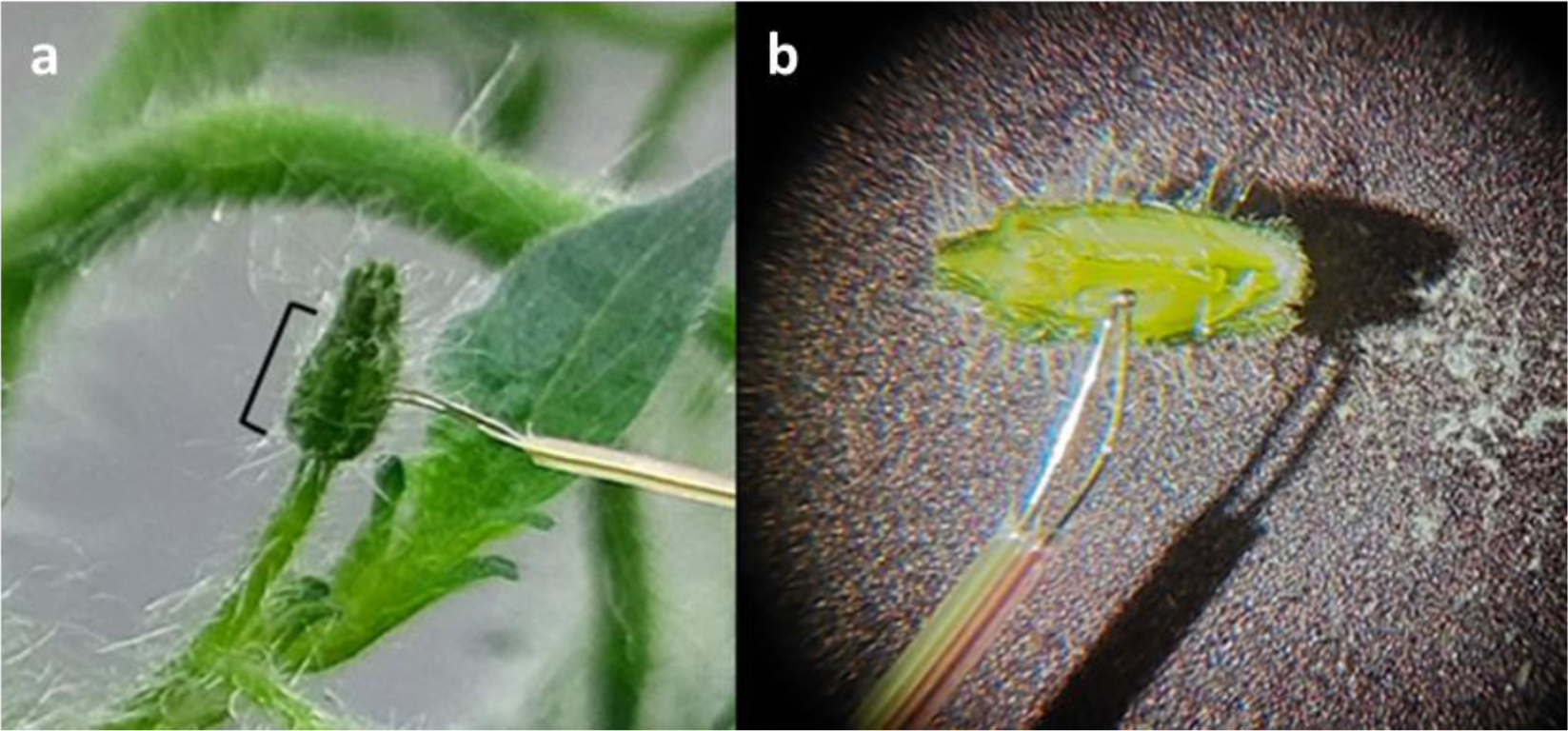
Measurement of bud and air temperature by thermocouple. To measure the flower bud core temperature, thermocouple type K sensors are inserted in flower buds of 3-4 mm. The sensor head is inserted halfway the cone shaped section of the flower bud, as indicated by black brackets (a) and pushed in until two pressure points are surpassed. At this moment the sensor head is present in the space between the pistil and anther cone in the center of the flower bud (b). Simultaneous with the measurement of the core flower bud temperature, a second thermocouple sensor connected to the same logger measures the air temperature at the same height in the vicinity of the flower bud.

**Extended data figure 2:**
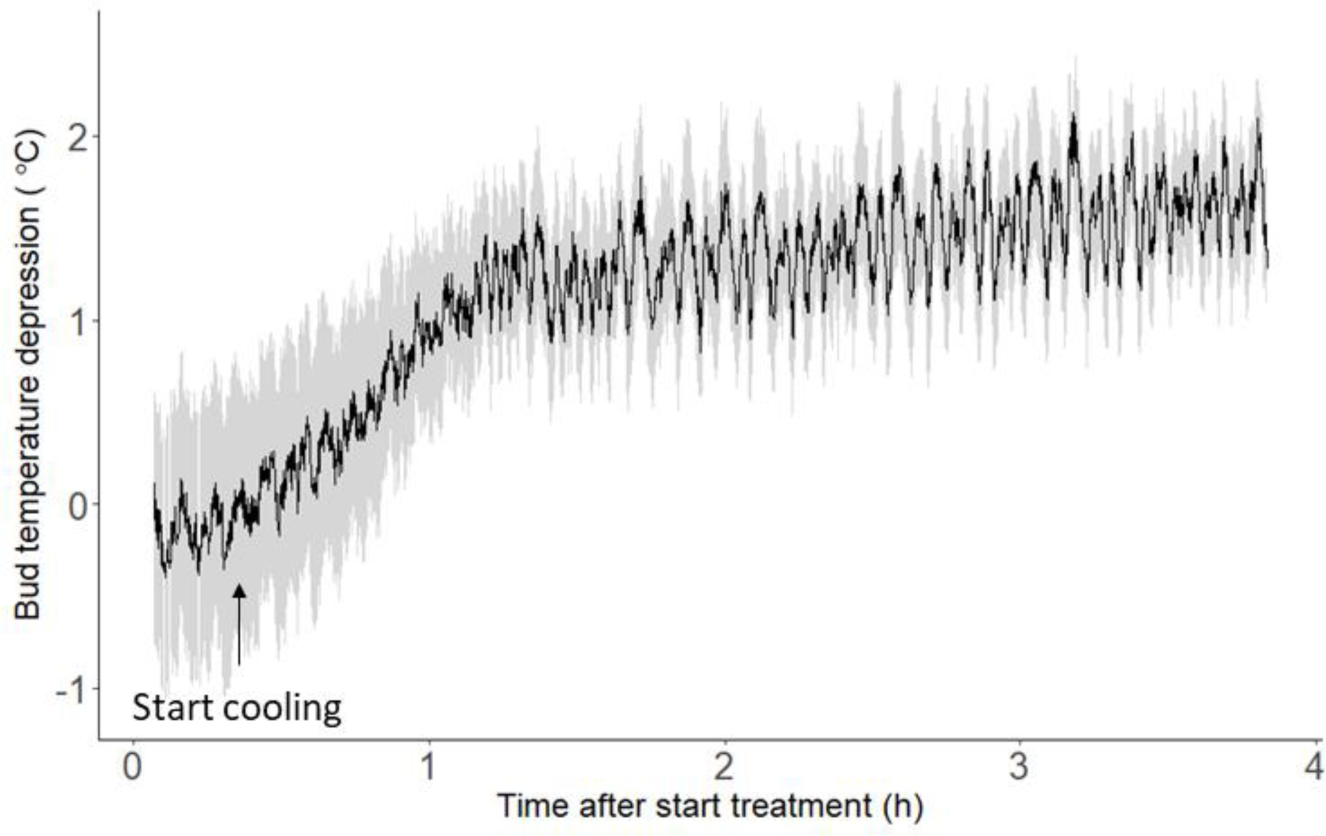
The bud temperature depression increases during the initial period of heat treatment. After a short period in the heat flower bud cooling is induced and the bud temperature depression slowly increases until it stabilizes at an air temperature dependent level. Indicated is the mean bud temperature depression and 95% confidence interval of flower buds (n=9 plants) upon placement in LTMH (34 °C).

**Extended data figure 3:**
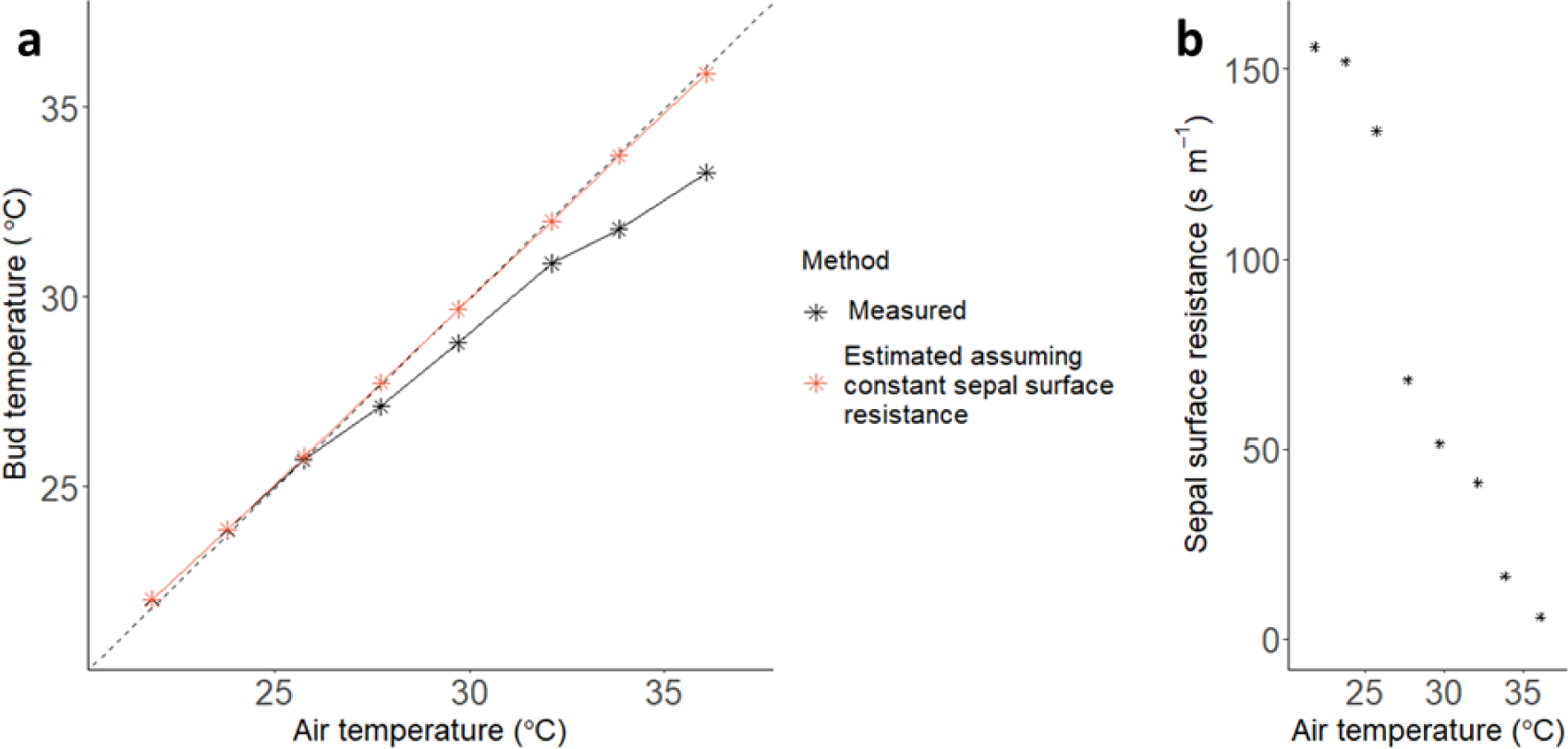
The sepal surface resistance decreases under increasing air temperature. To test if flower bud cooling required active regulation of stomata, the observed bud temperatures at air temperatures between 22 and 36 °C (a: black asterisks) were included in the energy balance, describing plant tissue thermoregulation^30^. These were used to calculate the sepal surface resistance at each temperature (b). The impact of this reduction in sepal surface resistance was subsequently visualized by estimating the bud temperature from the air temperature under the assumption of a constant sepal surface resistance / stomatal conductance (a: red asterisks), using the sepal surface resistance found at an air temperature of 22 °C. The line of equality (y=x) is indicated with a dashed line.

**Extended data figure 4:**
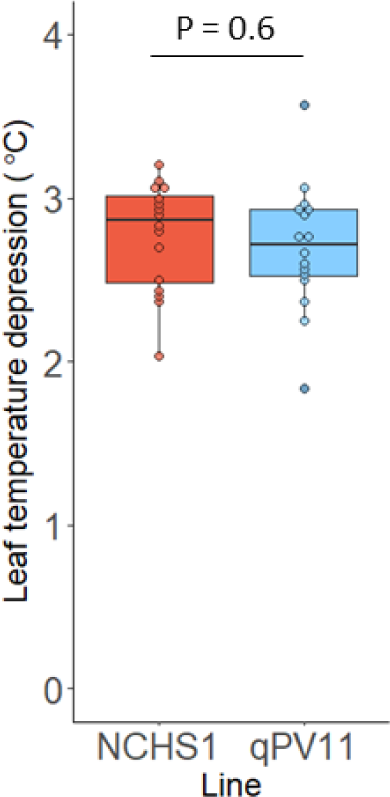
qPV11 does not affect leaf cooling. The effect of introgression of qPV11 on leaf cooling on the 5^th^ day of LTMH. Indicated are median and interquartile ranges of mean leaf temperature depression (n = 16 plants) of a NIL containing qPV11 and its respective background NCHS-1. N.s. Significance of differences was determined by two-way ANOVA including block effects.

## Methods

### Plant material, cultivation and heat treatment

Tomato (*Solanum lycopersicum*) cv. Micro-Tom^48^ (obtained from the National BioResource Project, Japan; accession TOMJPF00001) was used unless otherwise defined. The Micro-Tom introgression of *sitiens*^48–50^ was kindly provided by Dr Lázaro Peres (University of Sao Paulo, Brasil). Tomato cultivars Nagcarlang (LA2661) and NCHS-1 (LA3847) were obtained from TGRC and used to produce an F2 population^37^. The qPV11 (allele derived from cv. Nagcarlang) NIL was obtained by five-fold back-crossing of a selected F2 plant with NCHS-1 followed by selfing. The presence of the qPV11 allele was selected for on the region SL4.0ch11: 3806144…4039701 (∼1 cM). Plant cultivation and treatment was done as described previously for Micro-Tom^51^ (LTMH: 34/28 °C, 70% RH, 12/12h photoperiod) and qPV11/NCHS-1^37^ (LTMH: 32/26 °C, 70/80% RH, 14/10h photoperiod). For the reversed day-night cycle, temperature and corresponding relative humidity were switched between day and night. For other temperature settings, relative humidity was adjusted to maintain a vapor pressure deficit (VPD) equal to LTMH day settings. Flower bud and leaf temperatures were determined on the fifth day and pollen viability was determined during the third week of LTMH.

### Measurement of flower bud, leaf and air temperature and temperature depressions

Flower bud and air temperatures were measured every 5 seconds using a 6-channel temperature logger (DXL6SD-USB, OMEGA Engineering Inc.) with Ø 0.13 mm K-Type Thermocouple sensors (TL0260, PerfectPrime, London, United Kingdom). Flower bud temperature was measured by insertion of the thermocouple sensor head into the bud core, between the style and the anther cone of free-hanging flower buds of 3-4 mm (Extended data Fig. 1). Air temperature was measured simultaneously and at the same height with a second thermocouple sensor on the same logger. Air temperature was always measured by 3 air sensors, correlating to the 3 loggers used, unless otherwise indicated in figure legends. Temperature measurements started 50 minutes after insertion and logging continued for at least 25 minutes. New buds were measured daily in case of multiple-day logging. The same sensor was used in each contrasting group at least once in case two different treatments were compared. Temperatures were logged over series of entire cycles and were adjusted for sensor offsets per experiment based on simultaneous measurements of all sensors in still-standing air.

Flower bud and leaf temperatures of NCHS-1, qPV11 and sitiens were measured using a handheld thermocouple system (DEM106, Velleman, Gavere, Belgium) using the same K-type thermocouple sensors and methodology as described above. Leaf temperatures were measured by attaching the sensor head to the adaxial surface of leaves at similar height as measured flower buds using a clip-on. Air temperature was measured using a second sensor on the thermocouple system at equal height. Air and flower bud/leaf temperature were measured simultaneously and recorded when both temperatures were stable. Measurements close to the extremes of the thermo-amplitude of the climate room were avoided. Measured air temperatures were adjusted for sensor offsets per measurement based on simultaneous measurements of both sensors in still standing air and on a homogenous object.

Mean temperatures per plant per treatment/time period were used for boxplots and statistical analysis. Temperature depression is the difference between the air and tissue temperature and was averaged over the whole measurement period when used for boxplots and statistical analysis.

### Measurement of bud temperature depression at various air temperatures

To measure the bud temperature depression (BTD) at different air temperatures, air temperature was increased incrementally with 2 °C every 90 minutes on the fifth day of LTMH. Per temperature step, BTD was allowed to stabilize for 50 minutes and mean BTD was calculated over ∼30 minutes as described above. Sensors were installed the night before measurement and kept in place during the whole experiment. Measured bud temperatures for the various air temperatures were also used to determine changes in the bud surface resistance.

### Estimation of bud temperature from the heat energy balance

The observed air and flower bud temperatures were included in the heat energy balance^30^,

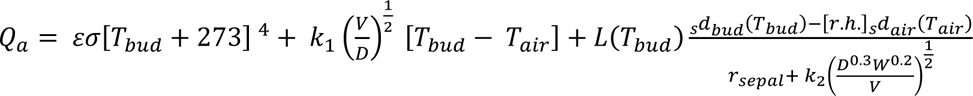

where Q_a_ is the total amount of radiation absorbed (assumed at 600 W m^-2^), ε is the emissivity of the sepal surface to longwave radiation (assumed at 0.96^52^), σ is the Stefan-Boltzmann constant, k_1_ is an empirical coefficient for the amount of energy required to heat the flower bud by 1 °C (assumed at 9.14 J m^-2^ s^-1/2^ °C^-1^), k_2_ an empirical coefficient for the boundary layer resistance (assumed at 183 s^0.5^ m^-1^), L(T_bud_) is the latent heat of vaporization of water which was taken as a constant (2.43 * 10^6^ J kg^-1^), V is the windspeed (estimated at 1 m s^-1^ in the climate cabinet), D is the width of the flower bud (∼1.9 mm), W is the length of the flower bud (∼ 4 mm), T_bud_ and T_air_ are the respective bud and air temperature in °C, _s_d_bud_ and _s_d_air_ are the respective saturation vapor densities of the flower bud and air in kg m^-3^, r.h. the relative humidity as a decimal fraction, and r_sepal_ the resistance to water vapor diffusion through the sepal surface in s m^-1^ (both through stomatal and substomatal pores). Net radiation, wind speed, bud shape and bud size were kept constant in the experiment allowing the extraction of the sepal surface resistance / resistance to water vapor diffusion^53–55^.

The resistance to water vapor diffusion was calculated from the observed stable air and flower bud temperatures for each temperature step during the incremental increase in temperature. The predicted flower bud temperature was calculated from the energy balance based on the air temperature, the above-mentioned variables and the assumption of a constant stomatal conductance. For a constant stomatal conductance the resistance to water vapor diffusion acquired at a T_air_ of 22°C was used for all temperature increments.

### Pollen viability assay

Flowers at the anthesis stage were sampled on at least 3 different days during the 3^rd^ week of LTMH. On each sampling day, 1-3 flowers per plant were pooled into a sample, and viability was determined for ∼10.000 pollen per sample. Pollen viability was determined through impedance flow cytometry, using Ampha™ Z32 with a D-chip, AF6 buffer, and other settings as recommended for tomato by the manufacturer (Amphasys AG, Luzern, Switzerland)^56^. Obtained mean pollen viabilities per plant were used for boxplots and logit transformed for statistical analysis.

### Transpiration measurements

Leaf and flower bud transpiration was measured of free hanging leaves and flower as described for temperature measurements using the LI-COR 6400 gas exchange system (LI-COR Biosciences, Lincoln, NE, USA) equipped with a leaf cuvette covering 6 cm^2^ area. Measurements were performed at a temperature of 32 °C, external CO_2_ concentration of 400 μL L^-1^, light intensity of 190 µmol PAR m^-2^s^-1^ and relative humidity of 67-70 % RH. Whole or partial inflorescences with flower buds of 1-4 mm were inserted intact in the chamber while on the plant. All tissue within the leaf chamber was sampled and used to determine fresh weight and total surface area. To determine total bud surface area, flower buds were scanned using a flatbed scanner (Epson 11000xl, Epson, Nagano, Japan) and scans were analyzed using root morphology analyzing software (WinRhizo 2013e, Regent Instruments, Quebec, Canada). Mean transpiration per tissue-plant combination was used for boxplots and statistical analysis.

### Manipulation of gas exchange by ABA and Vaseline treatment

To prevent sepal gas exchange, petroleum jelly (Vaseline, Sigma Aldrich, St. Louis, MO, USA) was gently applied to Micro-Tom flower buds. Bud temperatures of treated and untreated flower buds were measured for 1 hour, starting 80 minutes after application. Sensors were interchanged between treatments in a second round on the same plants.

To close stomata, 50 µM abscisic acid (ABA; Sigma-Aldrich) in aqueous solution (0.01% ethanol (v/v), 0.1% Tween20 (v/v)) or Mock (0.01% ethanol (v/v), 0.1% Tween20 (v/v)) was gently applied by brush to Micro-Tom flower buds. Bud temperatures of ABA and mock-treated flower buds were measured for 1 hour, starting 1 hour after application. Sensors were interchanged between treatments in a second round with new plants.

### Manipulation of stomatal dynamics by ABA and fusicoccin treatment

To prevent stomatal dynamics, pedicels of NCHS-1 and qPV11 plants were soaked with 5 mM ABA (1% ethanol, 0.1% Tween20) 90 minutes before measurement or 10 µM fusicoccin (0.1% ethanol, 0.05% Tween20) was gently applied to the flower bud by brush 1 day before measurement. Flower bud transpiration was measured alternately of NCHS-1 and qPV11 between 11:00 and 15:00.

### Generation and genotyping of RIL populations

A determinate and an indeterminate four-parental RIL population were generated for phenotyping in the field. An F1 hybrid between cultivars Nagcarlang (LA2661) and NCHS-1 (LA3847) was crossed with the determinate commercial hybrid ‘Nunhems Delfo F1’ (BASF Vegetable Seeds, Haelen, The Netherlands) to produce 400 F1 seeds for the determinate RIL population or with the indeterminate commercial hybrid ‘E15B.50893.’ (Enza Zaden, Enkhuizen, The Netherlands) to produce 400 F1 seeds for the indeterminate RIL population. These lines were propagated by single seed descent during 5 (determinate) or 3 (indeterminate) generations to produce the populations used in this study. The determinate / indeterminate nature of both populations was controlled by fixation of the respective alleles of *SP* (Solyc06g074350)^57^ during population development. Disease resistances were maintained in the whole population by fixation of the resistance genes *Mi1-2* (Solyc06g008450)^58^, *ve1* (Solyc09g005090)^59^ and *i2* (Solyc11g071430)^60^ in the determinate population and *tm2* (Solyc09g018220)^61^, *Ty-1* (Solyc06g051190)^62^ and *Ty-5* (Solyc04g009810)^63^ in the indeterminate population. Additionally *j2* (Solyc12g038510)^64^ was fixed in the determinate population for ease of mechanical harvesting and the Nagcarlang allele of the thermotolerance QTL qPV11^37^ was fixed in 25% of each population using solcap_snp_21012.

DNA was isolated from leaf tissue of the determinate RIL population and the next generation was grown in a processing tomato field. The indeterminate population was grown in a greenhouse and DNA was isolated from leaf tissue of the phenotyped plants. The presence of the qPV11 allele from Nagcarlang versus the three other parental alleles was scored based on the markers solcap_snp_sl_21005 (determinate population) and solcap_snp_sl_21012 (indeterminate population)^65^. Both populations had segregated for qPV11 and heterozygous individuals were removed before data analysis. From the 341 determinate RILs and 723 indeterminate plants homozygous for their qPV11 allele respectively 55 RILs and 202 plants were homozygous for the Nagcarlang allele of qPV11.

### Evaluation of pollen viability and fruit set under LTMH conditions in the field

The determinate RIL population was grown in a processing tomato field in Valdetorres, Spain (lat 38.935139, long −6.044229) during the summer of 2020. Temperature and relative humidity were measured every 10 minutes at plant height at 3 different locations in the field using HOBO MX2301A data loggers in RSB3B solar radiation shields (Onset, Bourne, MA, USA). Details on environmental conditions and moments of sampling are indicated in Supplementary Figure 1. Plants were grown following standard cultivation practices in two blocks of 35 rows and 12 columns, augmented with the parental lines as high frequency replicated controls. Each block contained the full RIL population, providing two replicates per RIL. The placement of genotypes was optimized using the DiGGer design package^66^. Each replicate consisted of a plot of 12 plants of the same line. Pollen viability measurements and fruit set estimations were taken per plot.

For pollen viability measurements, pollen were extracted from flowers on the first day of anthesis by vibrating the flower for 10 s using a sonic toothbrush (EZS 5663, AEG, Frankfurt, Germany). Pollen viability measurements were performed during the period of 7-11 days after a heatwave like event^10^.

Extracted pollen of 3 flowers of 3 different plants were pooled and pollen viability was measured by impedance flow cytometry as described previously. Pollen measurements occurred over multiple days and were averaged per plot for statistics. Fruit set was estimated as a breeding score from 1 to 9 by a professional processing tomato breeder at the end of season. The breeding score was determined as an estimation of the relationship between the number of seed-bearing fruits and the number of pedicels on the plant as estimated by eye. All phenotyping occurred prior to genotyping.

### Evaluation of pollen viability and fruit set under LTMH conditions in the greenhouse

The indeterminate RIL population was grown on drip fertigation inert substrate bags in a greenhouse in Almeria, Spain (lat 36.806474, long −2.765879), during the summer of 2021. Temperature and relative humidity were measured every 10 minutes at plant height at 2 different locations in the greenhouse as described previously. Details on environmental conditions and moments of sampling are indicated in the Supplementary Figure 2. Plants were grown following standard cultivation practices in two blocks of 52 rows and 8 columns, augmented with the parental lines as high frequency replicated controls. Each block contained the full RIL population, providing two replicates per RIL. The placement of genotypes was optimized using the DiGGer design package^66^. Each replicate consisted of one plant. Pollen viability measurements and fruit set estimations were taken per plant. During the period of 7-11 days after a heatwave like event, whole flowers in the first day of anthesis were sampled and the pollen viability was determined as described previously. Per measurement 1-3 flowers per plant were pooled dependent on the presence of flowers and plants were measured 2-3 times during the sampling period. The mean pollen viability per plant was used for statistics. To determine fruit set, inflorescences with flowers undergoing anthesis were tagged in the week post sampling. The total number of flowers which had undergone anthesis post sampling per tagged inflorescence was determined from the number of pedicels. The number of seeded fruits on the pedicels was counted after ripening. The average fruit set per plant was established as the number of seeded fruits per tagged flower that had undergone anthesis.

### Statistics and reproducibility

Sample sizes, statistical tests used, and P values are stated in the figure legends and more details are included in the supplementary information. In summary, bud temperature depressions above zero were determined by one-tailed one-way t-test, statistical differences between two treatments, lines or tissues were assessed by a Student’s t-test, Welch’s t-test or Mann-Whitney test as appropriate. A two-way ANOVA with treatment block as additional variable was used to assess statistical differences between two treatments when tested in two treatment blocks separated in time. Statistical differences between groups separated by two variables were assessed using a two-way ANOVA followed by Tukey’s HSD post-hoc analysis. The assumptions of a normal distribution and equal variance between treatment groups were tested by Shapiro-Wilk normality test and Levene’s or F-test respectively. All statistical analyses were performed in the R environment^67^.

### Reporting summary

Further information on research design is available in the Nature Portfolio Reporting Summary linked to this article.

## Data availability

Source data are provided with this paper

## Code availability

No custom codes were generated for this study.

## Acknowledgements

We thank S. Carli, G. Blanco Avila and S. Gunes for their advice and fruit set evaluations, the staff of the Radboud University Experimental Garden and Genebank (Nijmegen, The Netherlands) for their assistance in plant cultivation, S. Wessels for his work in the development of flower bud temperature measurements and S. Vialet-Chabrand for discussions on plant transpiration and stomatal regulation. We acknowledge ENZA Zaden and BASF Vegetable Seeds for the development of the used populations and their support and assistance in the field and greenhouse screening experiments. This work was supported by the Dutch Topsector Horticulture and Starting Materials (TKI TU grant number TU-2018-012, to IR) and the Dutch Organization for Scientific Research (NWO grant number 867.15.011, to IR; NWO grant number 482.20.214, to IR).

## Author information

### Author Contributions

M.J.J. and I.R. conceived and designed the study. M.J.J. planned and performed the characterization of flower bud temperature, the energy balance model calculations and the characterization of qPV11. S.Y.J. planned and performed analyses of the *sitiens* line. M.J.J., E.J.W.V. and K.K. planned and performed experiments on stomatal regulation. M.J.J., W.V., F.F.M., T.M., C.d.K., F.A.v.E. and I.R. planned and performed the field and greenhouse screening. M.J.J., I.R. and E.J.W.V. interpreted the data. M.J.J. and I.R. wrote the manuscript. All authors contributed to the article and approved the submitted version.

