## Supplemental figures for "Flower bud cooling protects pollen development and improves fertility during heatwaves"

### Supplementary data

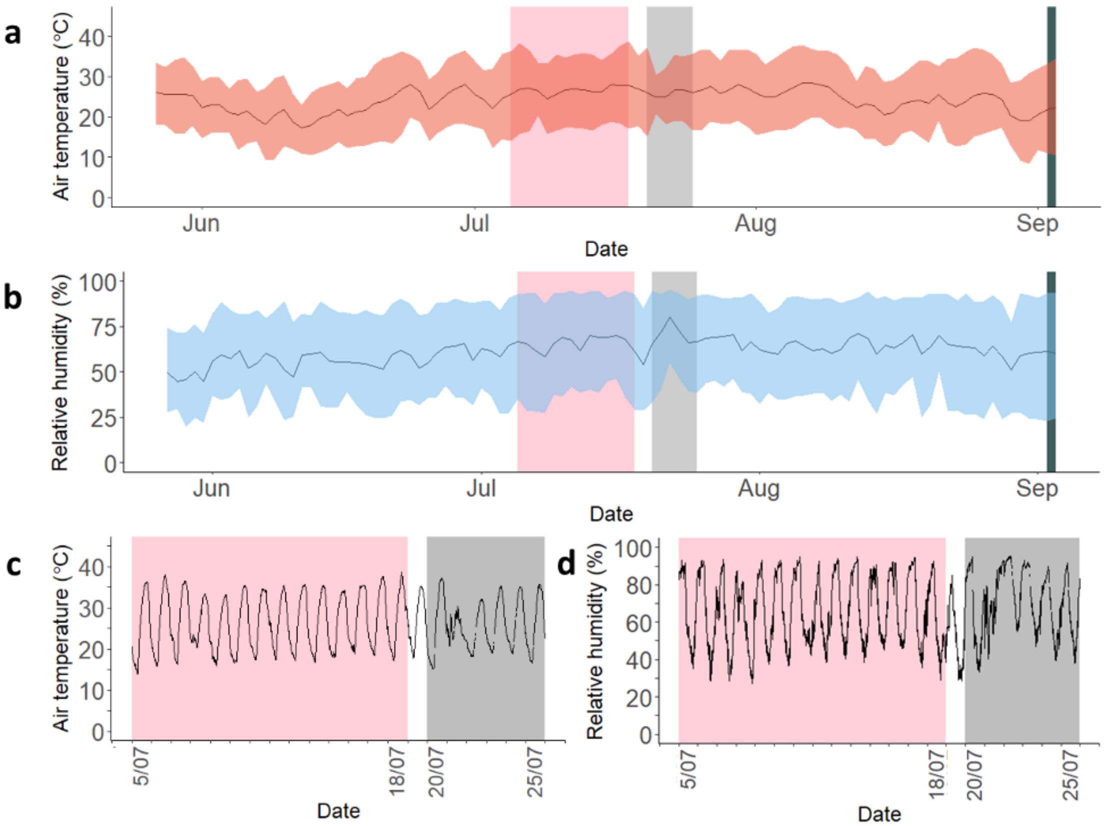


***Supplementary figure 1: Environmental conditions during the growth and pollen development of the determinate RIL population.*** *A determinate RIL population was grown on a processing tomato field in Valdetorres (Extremadura, Spain) during the summer of 2020. Air temperature (a) and relative humidity (b) were recorded during the entire growing season. Indicated are mean temperature / relative humidity (line) and maximum and minimum temperature range (ribbon) per 24 hours. For pollen viability, flowers in anthesis were sampled after a period of LTMH and fruit set was assessed at the end of the growing season. The period during which sampled flowers were in the sensitive stages of pollen development (pink), moment of sampling mature flowers for pollen viability assays (grey) and moment of fruit set assessment (dark grey) are indicated. Temperature (c) and relative humidity (d) data per 10 minutes are shown for the period of sampling and sensitive period of flower development of the sampled flowers.*

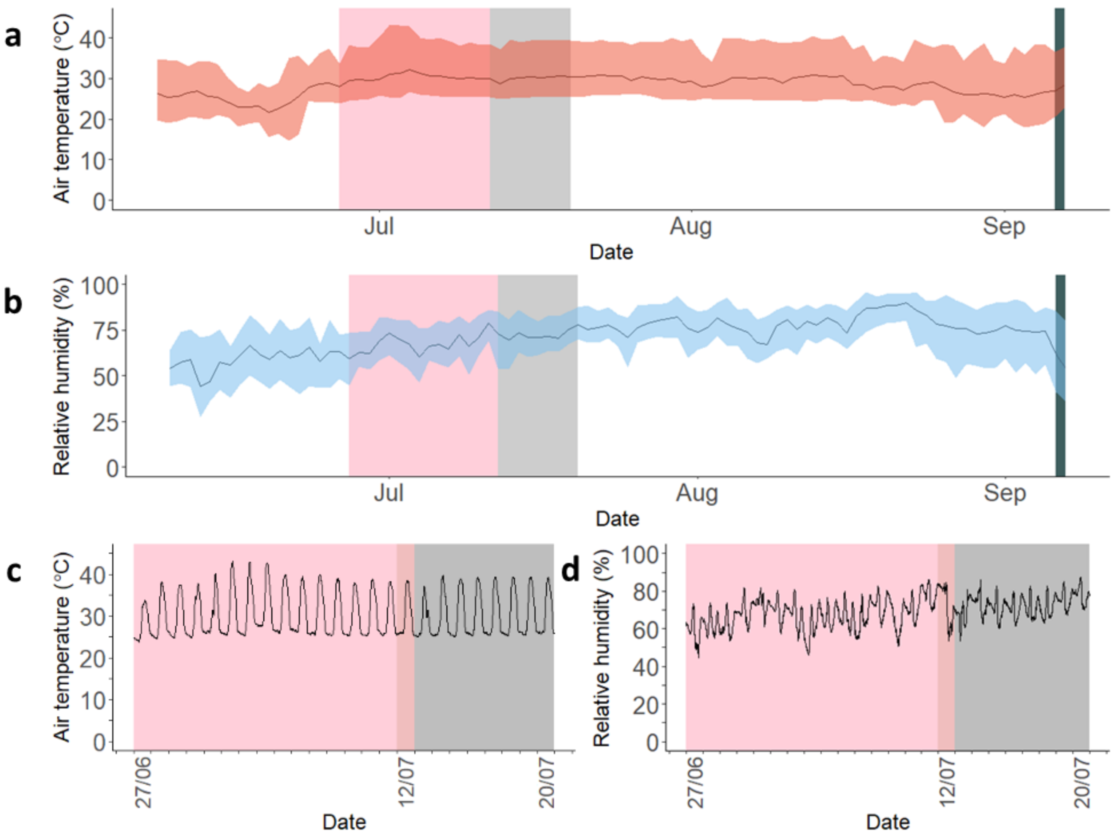


***Supplementary figure 2: Environmental conditions during the growth and pollen development of the indeterminate RIL population.*** *An indeterminate RIL population was grown in a greenhouse in Almeria (Spain) during the summer of 2021. Air temperature (a) and relative humidity (b) were recorded during the entire growing season. Indicated are mean temperature / relative humidity (line) and maximum and minimum temperature range (ribbon) per 24 hours. For pollen viability, flowers in anthesis were sampled after a period of LTMH and fruit set was assessed at the end of the growing season. The period during which sampled flowers were in the sensitive stages of pollen development (pink), moment of sampling mature flowers for pollen viability assays (grey) and moment of fruit set assessment (dark grey) are indicated. Temperature (c) and relative humidity (d) data per 10 minutes are shown for the period of sampling and sensitive period of flower development of the sampled flowers.*
